# Bacteria-powered living materials enable coral larval settlement

**DOI:** 10.1101/2024.12.18.629188

**Authors:** Natalie Levy, Samapti Kundu, Marnie Freckelton, Julie Dinasquet, Isabel Flores, Claudia T. Galindo-Martínez, Martin Tresguerres, Vanessa De La Garza, Yazhi Sun, Zahra Karimi, Crawford Drury, Christopher P. Jury, Josh Hancock, Shaochen Chen, Michael Hadfield, Daniel Wangpraseurt

**Affiliations:** Department of Chemical and NanoEngineering, University of California San Diego, La Jolla, CA, USA; Scripps Institution of Oceanography, University of California San Diego, San Diego, USA; Kewalo Marine Laboratory, University of Hawaiʻi, Honolulu, HI, USA 96813; Hawai‘i Institute of Marine Biology, University of Hawai‘i, Kāne‘ohe, HI, USA

**Keywords:** living materials, bacteria, bioprinting, ecosystem engineering, coral recruitment, coral reef restoration

## Abstract

The global decline of coral reefs calls for new strategies to rapidly restock coral populations and maintain ecosystem functions and services. Low recruitment success on degraded reefs hampers coral sexual propagation and contributes to limited genetic diversity and reef resilience. Here, we introduce a living bacteria-powered reef ink (Brink) for assisted coral recruitment. Brink can be rapidly applied to restoration substrates via photopolymerization, and it has been formulated to cultivate two settlement-inducing bacterial strains (*Cellulophaga lytica* and *Thalassotalea euphylliae*). Settlement assays performed with broadcast spawning (*Montipora capitata*) and brooding (*Pocillopora acuta*) Indo-Pacific corals showed that Brink-coated substrates increased settlement >5-fold compared to uncoated control substrates. Brink can be applied as a coating or 3D bioprinted, leading to various potential applications for integration with reef engineering. Our approach underscores the potential of using functional living materials for augmented ecosystem engineering and reef rehabilitation.

**Synopsis:** This study introduces a functional and sustainable bacteria-powered living material that enhances coral settlement, promoting coral reef rehabilitation and ecosystem resilience.

## 1. Introduction

Against the backdrop of the mounting pressures that global coral reefs face, such as rising ocean temperatures ^1^, ocean acidification ^2^, and pollution ^3^, replenishment of coral recruitment is vital to ensure their continued persistence. The restocking of reefs relies upon the successful recruitment of corals, ultimately building the reef habitat and maintaining biodiversity ^4,5^. Efforts in coral restoration typically involve either fragmenting adult corals or cultivating juvenile corals through sexual reproduction. Notably, the latter approach presents the advantage of bolstering genetic diversity ^6^, which is indispensable for enhancing ecosystem resilience ^7^. However, the progress of enhancing sexually propagated corals in their natural habitat has been hindered by low settlement rates ^8^. Therefore, there is an increasing interest in finding innovative ways to enhance coral recruitment in nature ^9^.

Coral recruitment relies on the sensory physiology of their larvae, which must detect and transduce chemical cues that stimulate settlement and metamorphosis ^10,11^. Some of the most well-documented attractants of invertebrate and coral larvae are chemical cues from marine bacterial biofilms ^12–18^ and crustose coralline algae (CCA) ^8,12,19–22^, an abundantly distributed functional group of algae that encrust the reef surface ^23,24^. More recently, individual strains of Gram-negative bacteria derived from biofilms, such as several strains of *Pseudoalteromonas* sp. ^12,13,16,19^ and *Cellulophaga lytica* ^15,25^, have been identified as invertebrate larvae morphogens ^12,13,16,19^. Bacteria from these genera produce conserved macromolecules, such as lipopolysaccharides (LPS) ^15,26^, that have been found to induce the settlement of invertebrate larvae, including marine worms, *Hydroides elegans* ^15^, and corals ^14,27^. The progress in understanding coral settlement induction opens avenues for interventions to enhance recruitment rates on reefs.

Using settlement-inducing bacteria to improve coral recruitment could be a viable strategy for reef restoration on natural, artificial, or hybrid reefs (i.e., man-made structures with biological interventions). Recent advances in biomanufacturing and 3D bioprinting have led to the development of functional living materials that facilitate the controlled growth of cells ^28^ and microorganisms ^29^. 3D bioprinting has revolutionized tissue engineering, precision medicine, and therapeutics ^30^. Recently, such techniques have been further developed for applications in environmental and marine sciences ^31^, including the engineering of microorganism-powered materials ^32,33^ that mimic coral tissues ^32^ and symbiosis ^29^ as well as functional synthetic biofilms ^34–36^. Controlling the microenvironment in which bacteria grow through the engineering of living materials, may lead to viable ways to enhance coral larval recruitment on natural and artificial coral reefs.

In the present study, we developed a living bacteria-powered reef ink (Brink) that enables the rapid biofabrication of hydrogels with settlement-inducing bacteria (Fig. 1). Hydrogel scaffolds were engineered to facilitate the stability and growth of two LPS-producing bacterial strains, *Cellulophaga lytica* and *Thalassotalea euphylliae*. Our study highlights the potential for developing functional living materials for coral rehabilitation and reef engineering ^31,37^.

**Figure 1.**
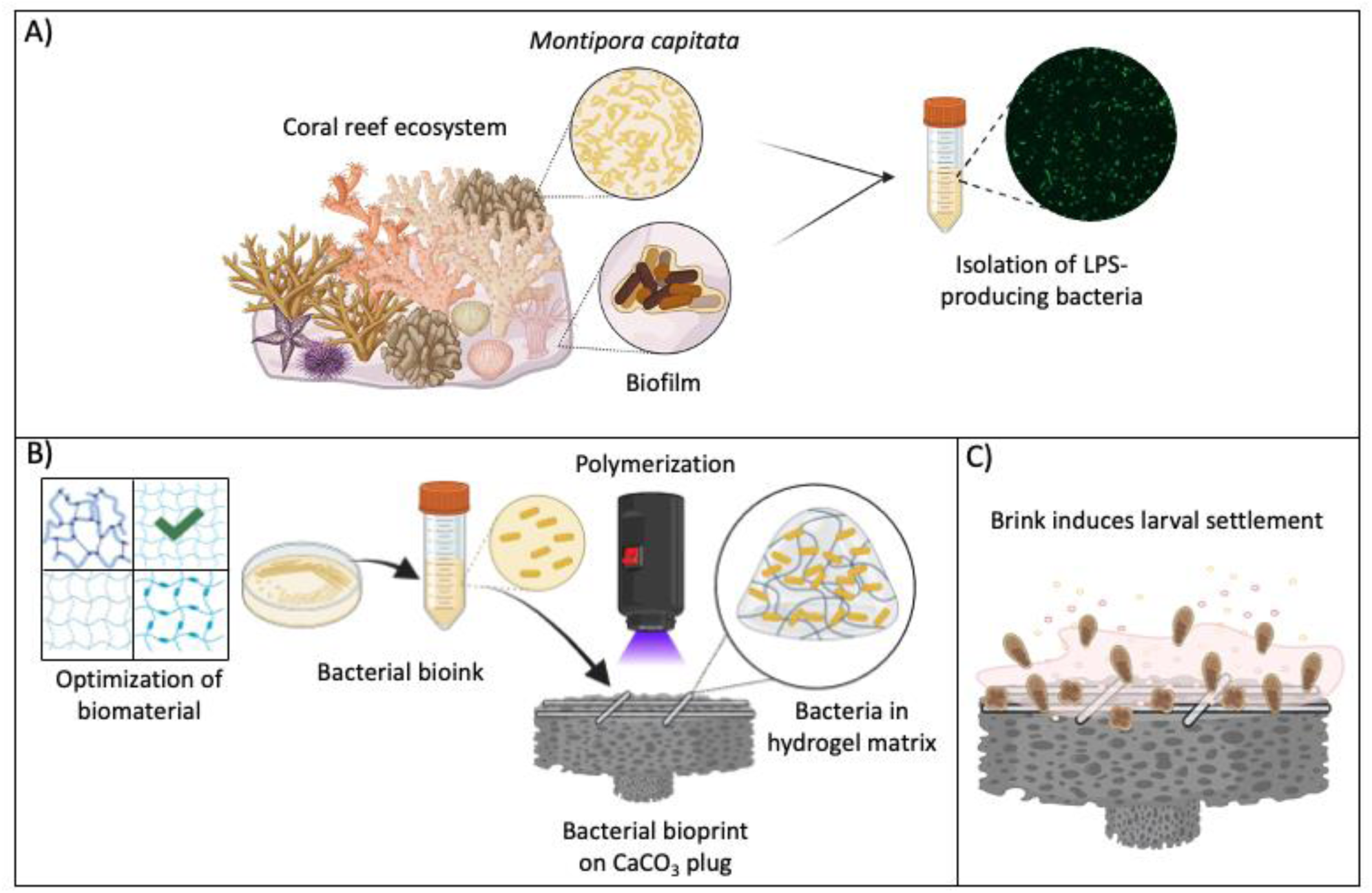
Overview of the engineering of functional living materials for coral settlement. **A)** Harvest and isolation of LPS-producing bacteria from the tissues of the coral *Montipora capitata* (*Thalassotalea euphylliae* H1) and marine biofilm (*Cellulophaga lytica* HI1). **B)** Optimization of biopolymer mixture for cell viability and mechanical properties. Light-assisted cross-linking of Brink on calcium carbonate (CaCO_3_) plugs, a common restoration material. **C)** The developed Brink hydrogel uses living bacteria as a bio-factory to produce chemical signals (LPS) and attract coral larvae from the environment.

## 2. Results and Discussion

### 2.1 Characterization of Brink coating as a viable microenvironment for bacteria

To develop Brink for rapid coating of reef substrates and coral settlement enhancement, we first identified suitable settlement-inducing bacteria. The bacterial strains, *T. euphylliae* isolated from coral tissues ^14,38^ and *C. lytica* isolated from marine biofilms ^18,39^, were selected based on their previously discovered ability to produce LPS and induce settlement of invertebrate larvae, including corals ^14,15,18,25–27^. To provide a tunable 3D cell culture environment for these bacterial strains, we optimized biopolymers based on high cell viability, biocompatibility, and long-term hydrogel stability. To tune their degradation, we utilized a combination of two known photocrosslinkable polymers frequently used in 3D bioprinting, poly(ethylene glycol)diacrylate (PEGDA) and gelatin methacrylate (GelMA) ^32^. PEGDA is a synthetic biopolymer that offers good mechanical stiffness, contributing to a higher degree of polymerization and anti-fouling properties ^40^. In contrast, GelMA is a natural biopolymer with high biocompatibility and biodegradability due to its low viscosity and highly porous structure ^40,41^. We developed a sturdy, yet porous hydrogel that provides a viable culture environment for settlement-inducing bacteria (Fig. 2).

**Figure 2.**
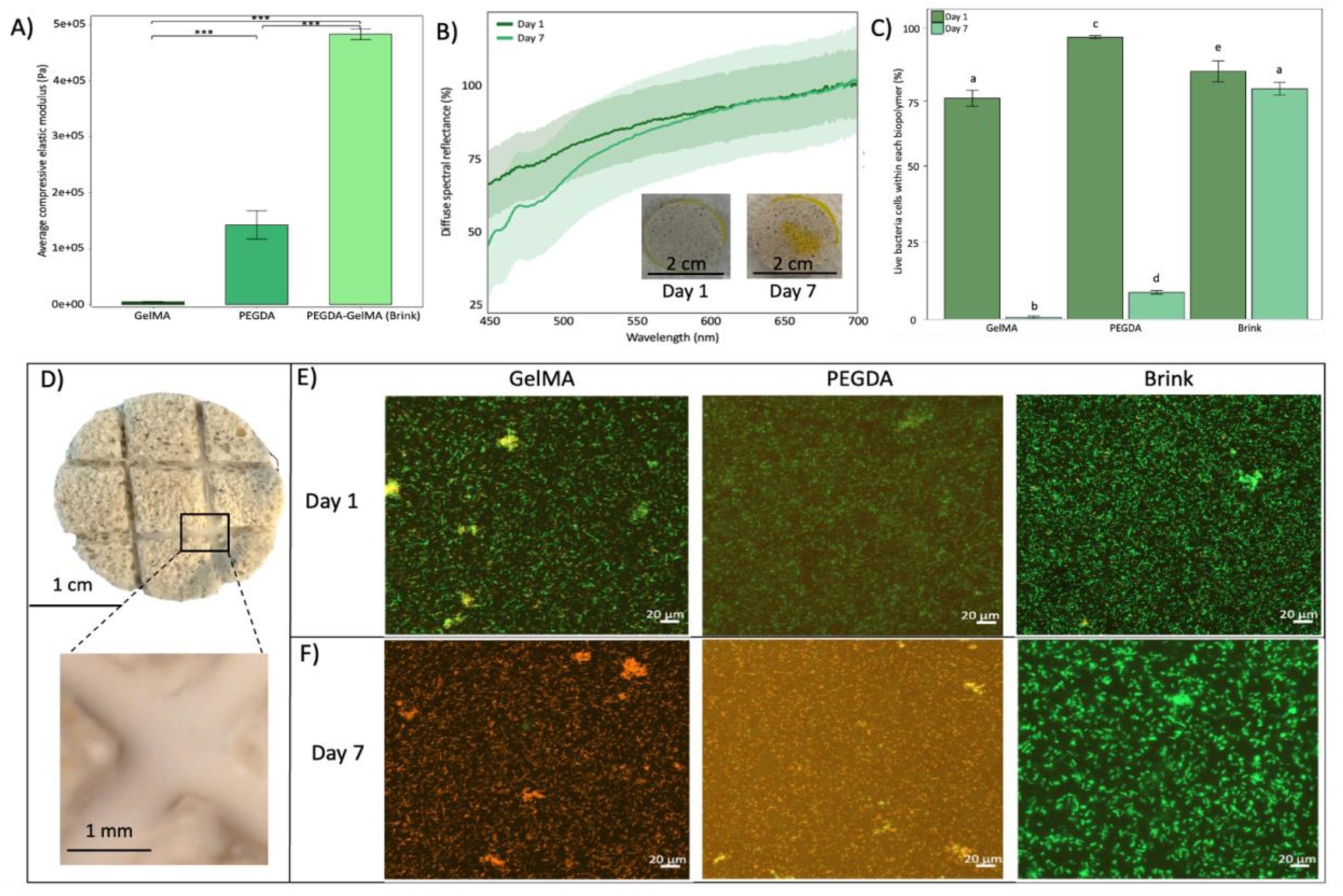
Characterization of the living bacteria reef ink (Brink). **A)** Average compressive elastic modulus ± SD (*n*= 3) of GelMA PEGA, and PEGDA-GelMA (Brink) hydrogels. Significance equivalent to *** p<0.0001. **B**) Diffuse spectral reflectance (%) of Brink hydrogels measured on day 1 and day 7. Shaded regions represent 95% confidence intervals (*n*= 5 hydrogels). Insets show the visual change in hydrogel transparency over seven days. **C**) Mean percentage of live cells ± SD at day 1 and day 7 (*n*= 3 hydrogels). Letters indicate significance among groups, with the same letters indicating no significance. **D)** Overview of CaCO_3_ plug with Brink hydrogel in crevices. **E)** Cell viability of *C. lytica* encapsulated in GelMA, PEGDA, and Brink hydrogels on day 1. Maximum z-projection images (orthogonal view, step size = 20 µm; stack volume 30 mm), showing live and dead cells, in green and red respectively, for GelMA, PEGDA, and Brink hydrogels on day 1. **F)** Live and dead cells (orthogonal view, step size = 20 µm; stack volume 30 mm) for GelMA, PEGDA, and Brink hydrogels on day 7 after one week.

Brink hydrogels are characterized by a high average Young’s modulus (Pa) (4.83 × 10^5^, SD ± 9.46 × 10^3^), which is about 110-fold higher than hydrogels that only contain GelMA (4.39E × 10^3^, SD ± 9.24E × 10^2^) and 3.4-fold higher than those that only contain PEGDA (1.42 × 10^5^, SD ± 2.5E × 10^4^) (ANOVA, p<0.0001, Fig. 2B). This structural rigidity of Brink hydrogels is essential for underwater applications, while its high porosity provides superior biocompatibility and cell affinity ^42,43^.

We tested bacterial growth and cell viability using *C. lytica*, by coating common calcium carbonate-based restoration substrate with GelMA, PEGDA, and Brink hydrogels (about 500 µm thick) and cultivated them for one week (Fig. 2B-C). Diffusive reflectance spectroscopy was used as a non-invasive tool to monitor cell growth and estimate bacterial density. The dense bacterial growth in Brink hydrogels from the day 1 to day 7 spectra was visible according to a reduction in diffuse reflectance (R_d_) between 450-600 nm ^44,45^. Likewise, bacterial growth was visually apparent according to the yellowing of the initially transparent Brink hydrogel (Fig. 2B, insets). Confocal imaging suggested that the proportion of dead cells was <12.2% (SD ± 10.7) at day one for all treatments and this includes any dead cells before photopolymerization (Fig. 2C). Thus, the short UV exposure (<45 s) and low UV intensity during photo-crosslinking had negligible impact on cell viability. After 7 days of cultivation in hydrogels, cell viability was low for GelMA (0.65%, SD ± 0.54) and PEGDA (9.5%, SD ± 0.64) hydrogels, while Brink-coated hydrogels sustained high cell viability of >81% (SD ± 2.23) (p<0.0001, respectively) (Fig. 2D-F), which is most likely due to the combination of PEGDA-GelMA (Brink) polymers allowing for a suitable, structurally sound and porous environment for the bacteria to thrive. The fabrication of Brink hydrogels thus provides a customizable microhabitat for the cultivation of settlement-inducing bacteria.

### 2.2 Induction of settlement of coral larvae with Brink

We tested the effect of Brink on larval settlement using the broadcast spawning coral, *M. capitata* and the brooding coral, *P. acuta* (Fig. 3A). Settlement of *M. capitata* (96 hours post-fertilization and 12-15 h after introduction to assays; Fig. 3B) was enhanced more than 5-fold on Brink-coated substrates compared to uncoated controls (p<0.0001). There was no significant difference in *M. capitata* settlement rates between *C. lytica* and *T. euphylliae* Brink-coated substrates. Over 82% of larvae settled either on the plug or the surface of the well in Brink treatments (Fig. 3C). Over 80% of settled *M. capitata* larvae settled directly on the plug and <20% settled on the surface of the well, regardless of bacterial treatment. The proportion of larvae that settled on the plug increased for Brink-coated substrates (80-88%, SD ± 2.96-3.668, p<0.05) compared to uncoated controls (70%, SD ± 3.85; Fig. 3D and Supplementary Figs. 1-2).

**Figure 3.**
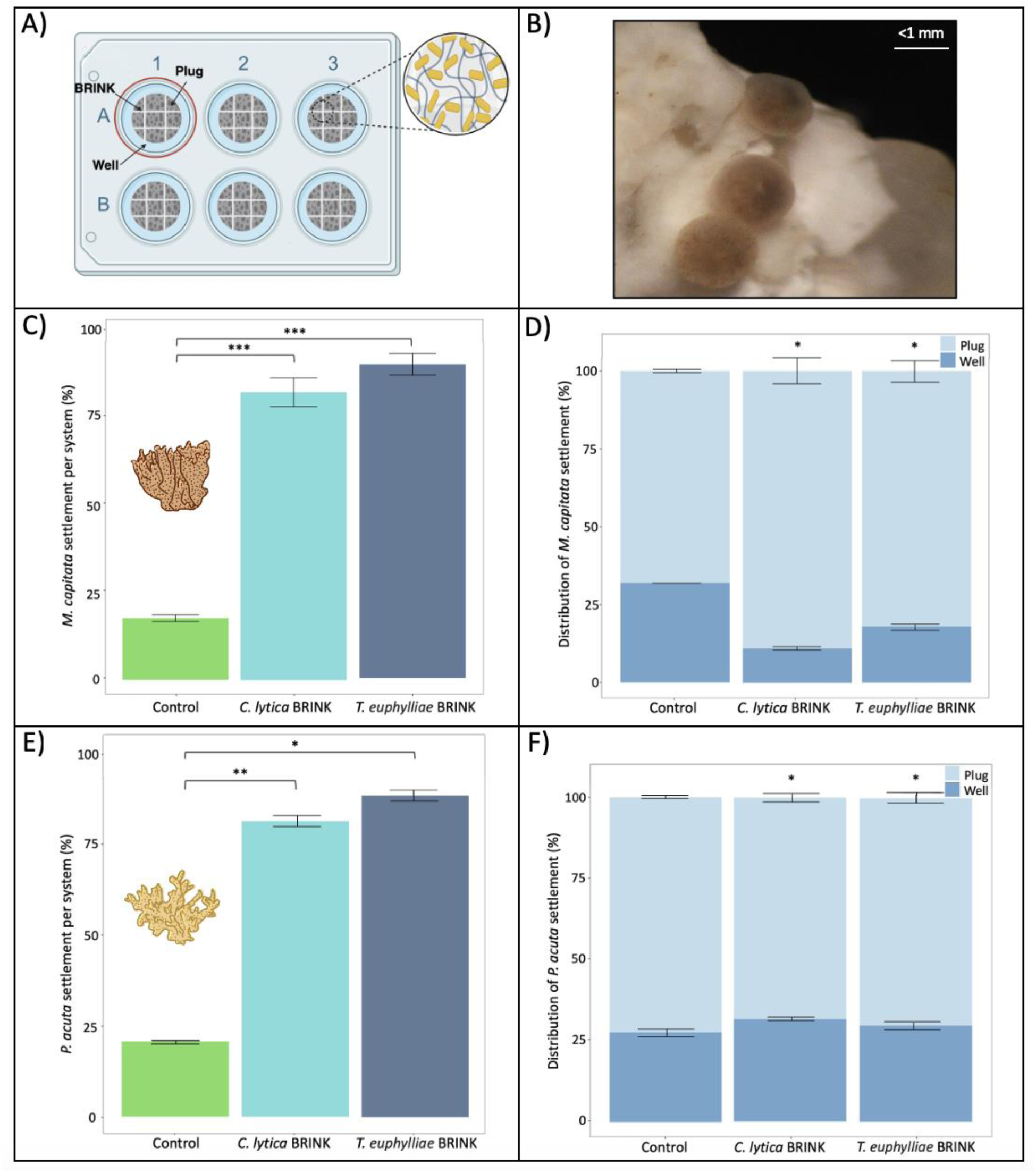
Settlement response of coral larvae to the engineered living Brink material. **A)** Experimental set-up of plugs in 6-well plates. Settlement was evaluated on the surface of the plug (Brink-coated or uncoated) as well as the surrounding surface of each well. **B)** Example image of settled and attached *M. capitata* larvae on a Brink-coated crevice. **C-D)** *M. capitata* settlement response. **C**) Total settlement (plug + well) in % of total larvae added (means ± SD, *n*= 6 wells). **D)** Distribution of settled larvae on the surface of the plug and the surrounding well-plate (means ± SD*, n*= 6 wells or plugs). **E-F)** *P. acuta* settlement response. **E**) Total settlement (plug + well) in % of total larvae added (means ± SD, *n*= 6 wells). **F)** Distribution of settled larvae on the surface of the plug and the surrounding well-plate (means ± SD*, n*= 6 wells or plugs). Significance levels are equivalent to * p<0.05, ** p<0.001, and *** p<0.0001. See also Figs. 1-4.

*P. acuta* larval settlement was enhanced more than 4-fold on Brink-coatings with over 81% settlement compared to uncoated controls (*T. euphylliae* 88.3%, SD ± 1.50, p<0.05; *C. lytica* 81%, SD ± 1.46, p<0.001; uncoated control 20.6%, SD ± 0.48; Fig. 3E and Supplementary Fig. 3). Similar to *M. capitata* larvae, there was no significant difference in *P. acuta* settlement rates between *C. lytica* and *T. euphylliae* Brink-coated substrates. Approximately 70% of the tested larvae preferred to settle directly on the Brink-coated plugs, while around 30% settled in the surrounding well (p<0.05; Fig. 3F and Supplementary Fig. 4). In contrast, only ∼13% of larvae settled on the plug of the uncoated substrates compared to the surrounding well (SD ± 0.38; Fig. 3F). These results suggest that the larvae settled in response to close proximal contact with the Brink coating. Although microbially mediated settlement induction is likely to vary for different coral species ^14,27^, the two bacterial strains used in this study were previously documented to induce settlement and metamorphosis in a wide range of marine invertebrates and could thus have settlement-enhancing effects on various coral species ^14,15,18,25–27,46^.

### 2.3 Brink microbial community dynamics in seawater

To assess the suitability of Brink for coral reef applications, we tested the bacterial community stability of Brink hydrogels in natural seawater for one week (Fig. 4). On day 3, *Cellulophaga* was the most abundant genus associated with Brink-coated plugs, indicating successful growth within the hydrogel matrix, despite the presence of opportunistic bacteria in the surrounding seawater (Fig. 4, Supplementary Fig. 5). Notably, *Cellulophaga* was not observed outside the hydrogel, thus suggesting effective immobilization with the hydrogel matrix. In contrast, the surrounding seawater was dominated by different bacterial genera over time, with eight genera prevailing on day 3 and another 10 genera persisting after a water change through day 7 (Fig. 4). A similar bacterial community was found on both Brink-coated and control hydrogels, suggesting that bacteria were introduced either via seawater or the polymer materials. By day 7, *Cellulophaga* abundance slightly declined, likely due to initial rapid growth and nutrient depletion (Fig. 4).

**Figure 4.**
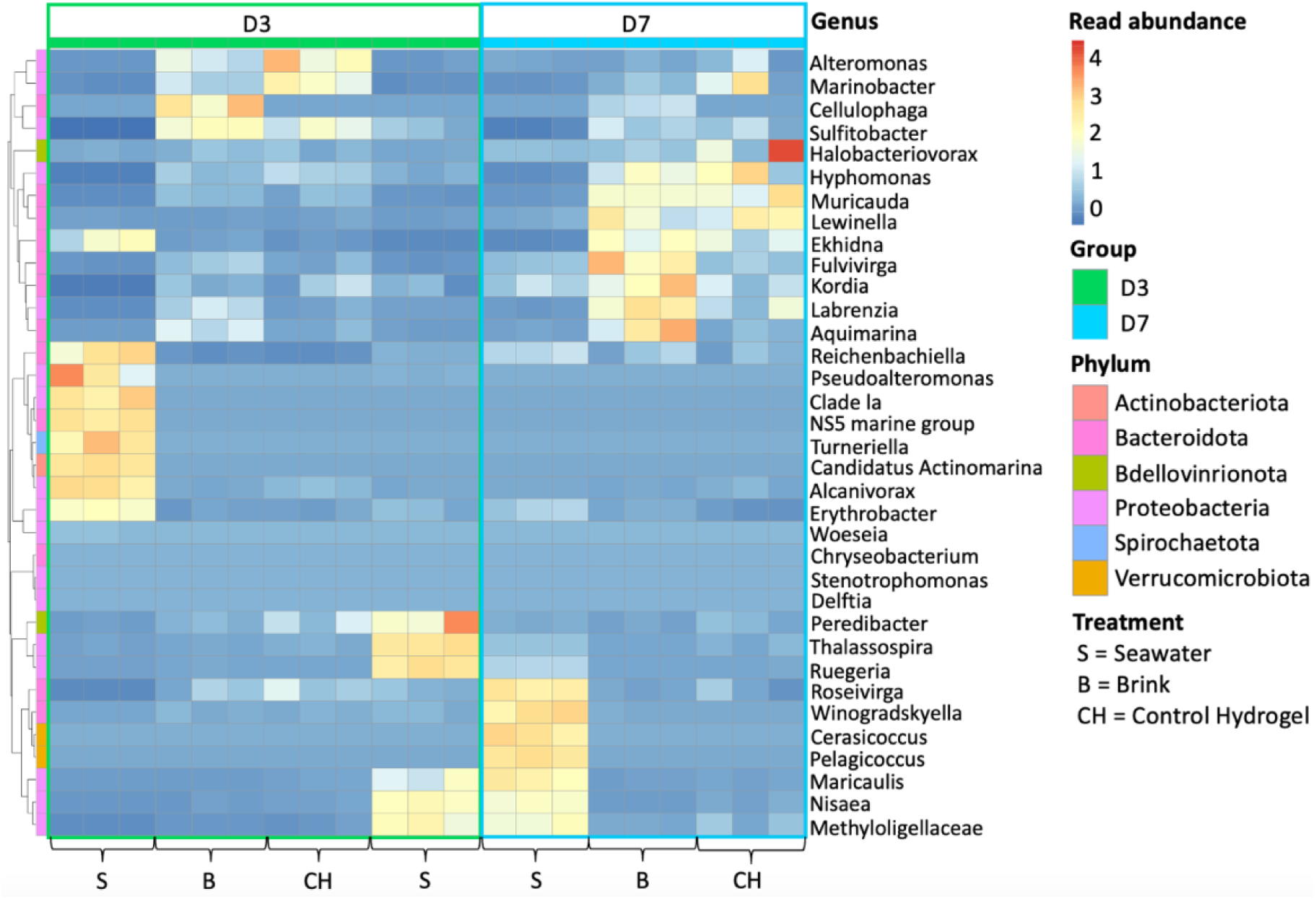
Normalized read abundances of the top 35 bacterial genera from 6 phyla associated with Seawater (S), Brink-coated plugs (B), and Control hydrogel plugs (CH) at day 3 (D3) and day 7 (D7) post-encapsulation (*n*= 3 per treatment). The first set of seawater replicates at D3 represents day 3 seawater, while the second set represents seawater after the day 3 water change. Scale bars and cell colors indicate the distance from the raw score to the mean standard deviation, with genera clustered by similarity.

## 3. Conclusions

In this study, we developed Brink, a bacteria-powered living material that significantly enhanced coral settlement in laboratory experiments, increasing settlement of *M. capitata* and *P. damicornis* by up to 5-fold compared to control substrates. The successful immobilization of bacteria within the hydrogel matrix in natural seawater suggests that Brink is suitable for *in situ* applications. For large scale applications in reef restoration and engineering projects, material synthesis and application must be done at scale. The biopolymers used in this study are either commercially available or can be easily produced, and bacteria cultivation at scale is well-established in industrial microbiology, providing a feasible pathway for large-scale production of Brink. Furthermore, light-assisted cross-linking is among the fastest methods for biomaterial processing, supporting its scalability. We envision that this bacteria-powered living material could be used to create effective settlement-inducing coatings for hybrid or biomimetic reefs ^47^, or 3D-printed settlement substrates ^48,49^ to accelerate the global efforts of restocking coral reefs.

## Materials and Methods

### Bacterial strains and culture

Individual strains of the Gram-negative bacteria *Cellulophaga lytica* (HI1) isolated from biofilms from a seawater table at Kewalo Marine Laboratory, Honolulu, Hawaii ^18,39^ and *Thalassotalea euphylliae* (H1) collected from *Montipora capitata* coral tissues ^14,38^ were obtained from the Kewalo Marine Laboratory (University of Hawai’i, Oahu, Hawai’i). The two bacterial strains were streaked from -80℃ glycerol stocks onto ½ filtered seawater (FSW) tryptone (½ FSWt) agar plates and placed into a 25℃ incubator for 24-48 h and kept in the dark ^15,25^. After incubation, 5 mL liquid bacterial cultures were prepared from ½ FSWt media and incubated for an additional 24 h at 25℃ on a shaker at 170 rpm. The liquid bacterial cultures were gently spun down (4,000 *g*, 30 min, 4°C) to collect bacterial pellets that were resuspended in 10 mL of sterilized FSW to achieve an optical density of 1.00 at OD_600_ with a cell density of ∼10^8^ cells ml^−1^ for both strains.

### Living bacteria reef ink (Brink) fabrication

To develop a suitable living bacteria-loaded ink, we tested and optimized a variety of photo-crosslinkable biopolymers. Biopolymers were chosen based on their known cell viability, mechanical stiffness, degradation properties, and ability to produce large batches for scalability. The combination of PEGDA and GelMA was chosen for their complementary mechanical properties ^40^. A combination of a higher percentage of PEGDA and a lower percentage of GelMA provides high resistance and minimal degradation of the hydrogel, which is a consequence of denser network structure providing higher crosslinking densities ^50,51^.

Gelatin methacrylate (GelMA) was synthesized through a direct reaction of porcine gelatin (Sigma Aldrich, St. Louis, MO, USA) and methacrylic anhydride (MA; Sigma Aldrich) as reported previously ^32^. Poly (ethylene) glycol diacrylate (PEGDA, Mn = 700 Da, Millipore-Sigma St. Louis, MO) and the photoinitiator lithium phenyl-2,4,6 trimethylbenzoylphosphinate (LAP) (TCI America™) were purchased. GelMA stock solutions (w/v) of 15% and 22.5% were prepared by dissolving lyophilized GelMA in MilliQ water. Brink was created by PEGDA, GelMA, LAP and bacterial solution at a final concentration of 10% PEGDA, 7.5 % GelMA, 0.5% LAP and 10^8^ cells ml^−1^ of bacteria (*C.lytica* or *T. euphylliae*). The bioink was protected from light and kept at room temperature until polymerization.

### Rapid free radical photopolymerization of the living bacteria reef ink

Calcium carbonate coral plugs (Ocean Wonders™) were chosen as the substrate for the hydrogels due to its composition, porosity, neutral pH, texture, and rugosity that makes it highly compatible with corals and their larvae. The plugs were dried in a 65℃ oven, sanded, engraved on top with 1 mm crevices (Fig. 1), rinsed, and dried again before use. 120 µL of Brink was polymerized onto the plugs. Photopolymerization was induced via a UV light source (405 nm, Thorlabs, New Jersey, USA) at an irradiance intensity of 17 mW/cm^−2^, which was optimized to facilitate successful cross-linking while minimizing potential cell damage^32^.

### Larval settlement assays

We performed settlement assays with a broadcast spawning coral species (*Montipora capitata*) and a brooding species (*Pocillopora acuta*) from O’ahu, Hawai‘i. For *P. acuta*, larvae were collected directly from adult colonies and introduced to settlement trials. *M. capitata* embryos were fertilized and reared to 96 hours post-fertilization following established best practices, before settlement experiments (Supplemental Materials).

Settlement assays were performed in 6-well plates containing 7 mL of FSW per well. We tested the efficacy of *C. lytica* Brink, *T. euphylliae* Brink, and uncoated control plugs. For all experiments, plugs were placed in 6-well plates upside down, with the top of the plug containing crevices face down in the well (*n*= 6 per treatment). Plugs were acclimated for a few hours in the well plates before adding coral larvae. Coral larvae were rinsed with FSW and counted to obtain an estimated number of larvae per mL. Due to the differences in size of the larvae, number of colonies producing larvae, and initial collection densities from each coral species, settlement experiments that involved *P. acuta* (∼500 µm - 1 mm larval length) used a density of ∼10-12 larvae per well for each treatment. Experiments with *M. capitata* (∼300 µm larval length) used 3 mL of larvae in FSW per well (∼40-45 per well), with a density of 15 larvae per mL for each treatment. Larvae were added to the wells and covered loosely with aluminum foil to facilitate oxygenation while preventing light exposure. Well plates were maintained overnight in a 27℃ incubator with a 12-12 h light-dark cycle at HIMB. After 12-15 hours, plugs were agitated to remove non-attached larvae and the number of attached, dead/disintegrated, and swimming larvae in each well was quantified using microscopy via a stereo microscope (Motic SMZ-168) with a camera attachment (Cannon EOS 600D) ^52^.

### Viability of living bacteria reef ink coating

Viability of *C. lytica* in the different polymers was assessed using the LIVE/DEAD™ BacLight™ Bacterial Viability Kit (Invitrogen, Thermo Fisher Scientific, Oregon, USA) and confocal microscopy. The kit was used following the manufacturer’s protocol with minor modifications that included 15 min incubation with the stain at room temperature in total darkness followed by rinsing the hydrogel gently with sterile FSW. The stained hydrogels were then visualized using an inverted confocal microscope AxioObserver Z1 with LSM800 and Zen 2.6 blue edition software (Carl Zeiss, Oberkochen, Germany), a 20x objective lens (Plan-Apochromat 20X/0.8 M27), and optimized settings for EGFP (to visualize live bacteria; excitation 480/500 nm at 0.1-0.15 % laser power, emission 488nm, detection 410-546 nm), and TuRFP (to visualize dead bacteria; excitation 490/635 nm at 0.45% laser power, emission 561nm, detection 400-650 nm). The Z-stacks (30 mm) were then presented as maximum intensity projection.

A calibration curve for reflectance was created by correlating the optical density (OD) of bacterial liquid cultures with the absorbance (*De*) in Brink, to estimate bacterial density. OD was first measured in 2 mL of concentrated bacterial culture, followed by a serial dilution, where 1 mL of the culture was replaced with fresh media. OD was measured after each dilution, and this process was repeated eight times. The removed bacterial culture from each step was used to polymerize the living Brink coating, following the photopolymerization method previously described, and De was calculated for each dilution. OD in the liquid bacterial cultures was determined using a miniature spectrometer (Flame, Ocean Optics, USA; *n=* 5 scans per measurement, boxcar width= 2 nm, resolution= 0.2 nm) in transmission mode. The liquid bacterial culture was placed into 3 mL cuvettes, and the absorption spectra were measured between 400 and 750 nm. Attenuation values were corrected by subtracting absorbance at 750 nm.

*De* spectra from Brink coatings were calculated from reflectance (*R*) measurements as *De* = Log(1/*R*) according to ^53–55^. Measurements were performed between 400 and 750 nm using a reflectance probe attached to a miniature spectrometer (Flame, Ocean Insight; *n=* 5 scans per measurement, boxcar width= 2 nm, resolution= 0.2 nm). The probe was placed 5 mm away from the Brink coatings’ surface at a 45° angle relative to the surface. Three random surface regions per bacterial coating were chosen for the measurements, and the experimental outcomes were normalized against a 99% diffuse reflectance standard (Spectralon, Labsphere, USA). Finally, bacterial growth in the Brink coating was monitored by determining bacterial *De* every day after polymerization for an entire week at 0 h, 12 h, 24 h, 48, 5 days, and 8 days after encapsulation of the bacteria.

For cell counts, samples (1 mL) for each isolate were fixed with 0.2 µm filtered formaldehyde (2% final concentration) for 15 min and stored at -80°C until processing. Bacterial cells were stained with SYBR Green I at 0.025% (vol/vol) final concentration, and counts were performed using a guava EasyCyte 5HT flow cytometer (Millipore©, USA).

Cell counts of the Brink coating were analyzed from confocal images in ImageJ2 Fiji (v.2.9.0/1.53t). Living and dead cells were quantified by creating a 16-bit black-and-white image. As the bacteria were rod-shaped, the cells were then rounded, the background noise was reduced, the image was converted to binary, and the cells were then individually separated. Cell sizes were determined by pixel units (100-1500), outlined for circularity, and edges were excluded.

### Mechanical stiffness

Mechanical properties were measured with a commercial device MicroSquisher (CellScale). Hydrogel cylinders of GelMA 22.5%, PEGDA 30%, and PEGDA-GelMA 30% and 15% with a height of 1 mm and a diameter of 1 mm, were printed using a custom-made 3D bioprinter (*n=* 3). The hydrogel scaffolds were printed with a cylindrical mask (1 mm x 1 mm) onto methacrylate glass coverslips (405nm for 30 s at 17 mW/cm^−2^), using PDMS spacers with a thickness of 1 mm. Samples were compressed by a cantilever at the speed of 3 μm/s to reach the strain rate of 18%, and were held for 2 s, then recovered to the original size at a speed of 6 μm/s. Each sample was compressed three times. The first two compression cycles removed hysteresis caused by internal friction. Young’s elastic modulus was calculated using an in-house MATLAB code with the force and displacement data of the third compression cycle from the CellScale software.

### Microbial community of Brink coatings in seawater

To understand the potential changes to the monoculture bacterium community (i.e., *C. lytica*) inside the hydrogel environment when exposed to natural seawater, an *ex-situ* aquarium experiment was performed at Scripps Institution of Oceanography (SIO). The aquarium set-up contained a heater set at 26°C, an air stone for oxygen, a powerhead for flow, and lighting set to a 12-12 h cycle (100 µmol/m^2^/s) to mimic natural conditions in a shallow reef ecosystem. Coral fragment plugs (Ocean Wonders™) were sanded, rinsed, and evenly coated with a top layer (∼2 cm in diameter with a ∼500 µm thickness) containing either a Brink hydrogel or a control hydrogel (hydrogel containing no bacteria). On day 1 (D1) plugs with the Brink hydrogel (*n=* 7) and control hydrogel (i.e., PEGDA-GelMA) (*n=* 7) coatings were polymerized with UV light and prepared according to the same protocol as described above. One plug from each treatment was immediately sampled by scraping the polymerized coating off the plug with a sterile scalpel blade and deposited into a 2 mL cryovial tube, which was immediately snap-frozen at -80°C prior to DNA extractions and 16S rRNA gene sequencing. Ambient 26°C seawater was filtered (0.45 µm) from the SIO pier to obtain a seawater community similar to remove any particles and potential grazers. Both plug treatments were spaced and distributed in an eggcrate sitting mid-level in the same aquarium (at a level of ∼90 µmol/m^2^/s). A 1 L seawater sample was collected from the aquarium before plugs were placed, which was then vacuum filtered onto a 0.2 µm filters (Pall Supor® 0.2 µm and 47 mm). After filtering the seawater three times, it was deposited into a cryovial tube and immediately snap-frozen at -80°C to preserve for further downstream analysis. Three plugs from each treatment were sampled again on day 3 (D3) and day 7 (D7). A 1 L seawater sample was collected on D3 before plugs were removed from the aquarium and filtered and preserved. Aquarium water was changed after the D3 plugs were removed and the D3 seawater samples were taken. New 26°C filtered seawater (0.45 µm) replaced the old water and a 1 L sample of the new seawater was collected, filtered, and preserved. The same seawater filtration and preservation process was repeated on D7 to mark the final bacteria community in the seawater at the end of the experiment.

DNA samples were extracted from Brink-coated coral fragment plugs, control hydrogel-coated plugs, and the seawater from the aquarium using the DNeasy® PowerBiofilm® Kit (Qiagen, Germany) according to the manufacturer’s protocol. PCR amplification was conducted to target the 16S rRNA gene V4-V5 region (450bp fragment length), using the universal primer set 515F and 907R (GTGCCAGCMGCCGCGGTAA and CCGTCAATTCCTTTGAGTTT, respectively)^56,57^. PCRs, amplicon purification, library preparation, amplicon sequencing, quality control, and bioinformatics were conducted by Novogene.

Raw reads were spliced and filtered to obtain clean data before using DADA2 to reduce noise ^58^. The remaining amplicon sequence variants (ASVs) were annotated to identify the related species information and abundance distribution ^58^. Reads were trimmed, filtered, denoised, merged, and chimeras were removed using the DADA2 workflow with QIIME2’s classify-sklearn algorithm ^59,60^, which is a Naïve Bayes classifier for species annotation of individual ASVs (trimmed to V4-5 region) using the 16S database SILVA v138.1 ^61^. The final ASVs were assigned to taxonomies and transferred from QIIME2 to R for statistical analyses.

### Statistical analyses

A one-way analysis of variance (ANOVA) and post-hoc tests were performed for mechanical stiffness (compressive elastic modulus) of each biopolymer hydrogel scaffold. Proportion data for live and dead cells in different biopolymers and larval settlement assays were tested for normality of variance with a Shapiro-Wilk test and homogeneity of variance with Levene’s test, where they failed to meet these assumptions^21^. The data was log-transformed, which did not affect the normality or homogeneity of variance, and a non-parametric analysis was conducted with the Kruskal-Wallis Rank Sum Test and the Dunn’s post-hoc test ^21^. Statistical significance was determined with p<0.05 and bar graphs were plotted as averages with ± SD. ASV counts from QIIME2 and metadata were imported into R to conduct 16S amplicon bacteria community analyses, producing a heatmap visualized via the pheatmap R package ^62^. All statistical analyses and plots were conducted using R (v4.0.3 and v4.2.1 ^63^.

## Acknowledgments

We acknowledge the support of the R3D consortium, specifically Ben Jones, Josh Levy, Josh Madin, Dan Schar, and Lindsey Badder for experimental and administrative support. Figures 1 and 3 were created using elements of BioRender (. The research work presented in this article is supported by the Defense Advanced Research Projects Agency under the Reefense Program. The views, opinions and/or findings expressed are those of the author and should not be interpreted as representing the official views or policies of the Department of Defense or the US Government.

## Author contributions

Designed the study: NL, DW. Performed the experiments: NL, SK, IF, CGM, VG, YS, ZK. Analyzed and interpreted the data: NL, MF, JD, IF, CGM, YZ, MH, DW. Contributed reagents/materials/tools/animals: MF, CD, JH, CPJ, SC, MT MH, DW. Wrote the main text of the manuscript: NL, with editorial contributions from all authors.

## Data availability

Sequence data from this study will be made available through a public repository.

## R3D consortium

Ben Jones¹, Josh Levy¹, Sean Mahaffey¹, Aricia Argyris¹, Mark Aruda¹, Ian Robertson¹, Zhenhua Huang², Ayrton Medina-Rodriguez², Mert Gokdepe², Brady Halvorson², Jon Chase², Charlotte White², Cami Dillon², Kristian McDonald², Anna Mikkelsen², Josh Madin³, Mollie Asbury³, Jessica Haver³, Hendrikje Jorissen³, Nina Schiettekatte³, Marion Chapeau³, Rob Toonen³, Chris Suchocki³, Van Wishingrad³, Christopher P. Jury³, Dan Schar³, Madeleine Hardt³, Claire Lewis³, Claire Bardin³, Joshua Kualani³, Crawford Drury³, Kira Hughes³, Josh Hancock³, Carlo Caruso³, Andrea Grottoli⁴, Shannon Dixon⁵, Josh Voss⁵, Allison Klein⁵, Sid Verma⁵, Alejandro Alvaro⁵, Richard Argall⁶, Kevin Chun⁶, William Hicks⁶, Alex LeBon⁶, John Yeh⁶, Aaron Thode⁷, Oceane Boulais⁷, Daniel Wangpraseurt⁷, Samapti Kundu⁷, Natalie Levy⁷, and Lindsey Badder⁷

^1^Applied Research Laboratory at the University of Hawaiʻi

^2^University of Hawaii at Manoa ^3^Hawai‘i Institute of Marine Biology ^4^Ohio State University

^5^Florida Atlantic University

^6^Makai Ocean Engineering

^7^Scripps Institution of Oceanography, UC San Diego

