## Supplemental Data for "Bacteria-powered living materials enable coral larval settlement"

Natalie Levy et al.

This PDF file includes:

Supplementary Figures 1-5
Supplementary Text
References 1-2

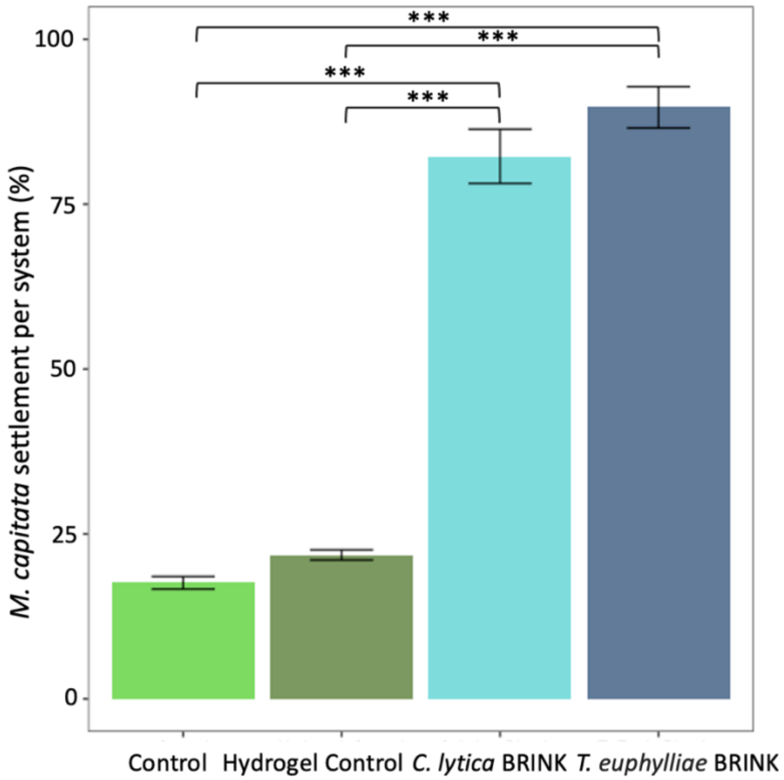


**Figure 1. Settlement response of *M. capitata* to the engineered living Brink hydrogel coating.** Total settlement (plug + well) in % of total larvae added (means ± SD, *n*= 6 wells). Significance levels are equivalent to * p<0.05, ** p<0.001, and *** p<0.0001.

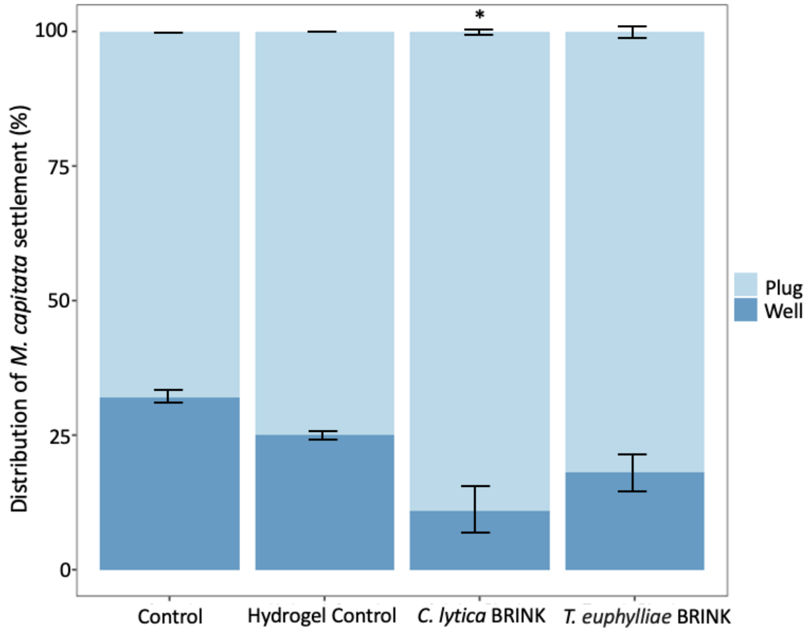


**Figure 2. Settlement response of *M. capitata* to the engineered living Brink hydrogel coating**. Distribution of settled larvae on the surface of the plug and the surrounding well-plate (means ± SD, *n*= 6 wells or plugs). Significance levels are equivalent to * p<0.05.

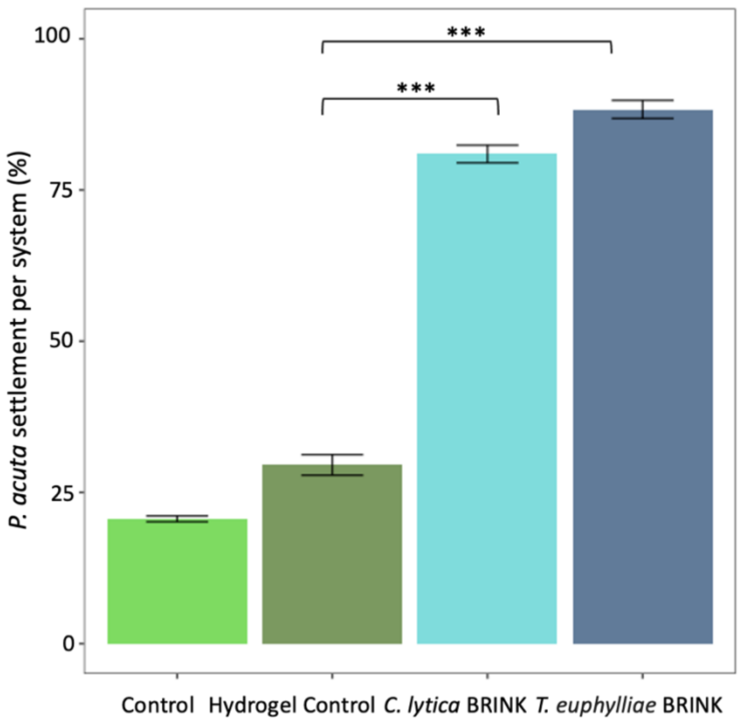


**Figure 3. Settlement response of *P. acuta* to the engineered living Brink hydrogel coating.** Total settlement (plug + well) in % of total larvae added (means ± SD, *n*= 6 wells). Significance levels are equivalent to * p<0.05, ** p<0.001, and *** p<0.0001.

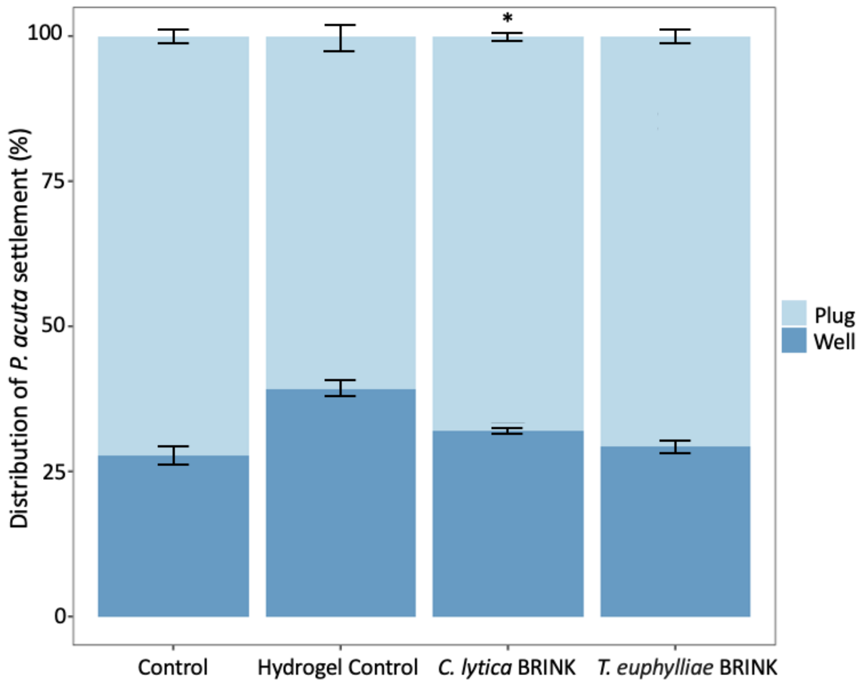


**Figure 4. Settlement response of *P. acuta* to the engineered living Brink hydrogel coating**. Distribution of settled larvae on the surface of the plug and the surrounding well-plate (means ± SD, *n*= 6 wells or plugs). Significance levels are equivalent to * p<0.05.

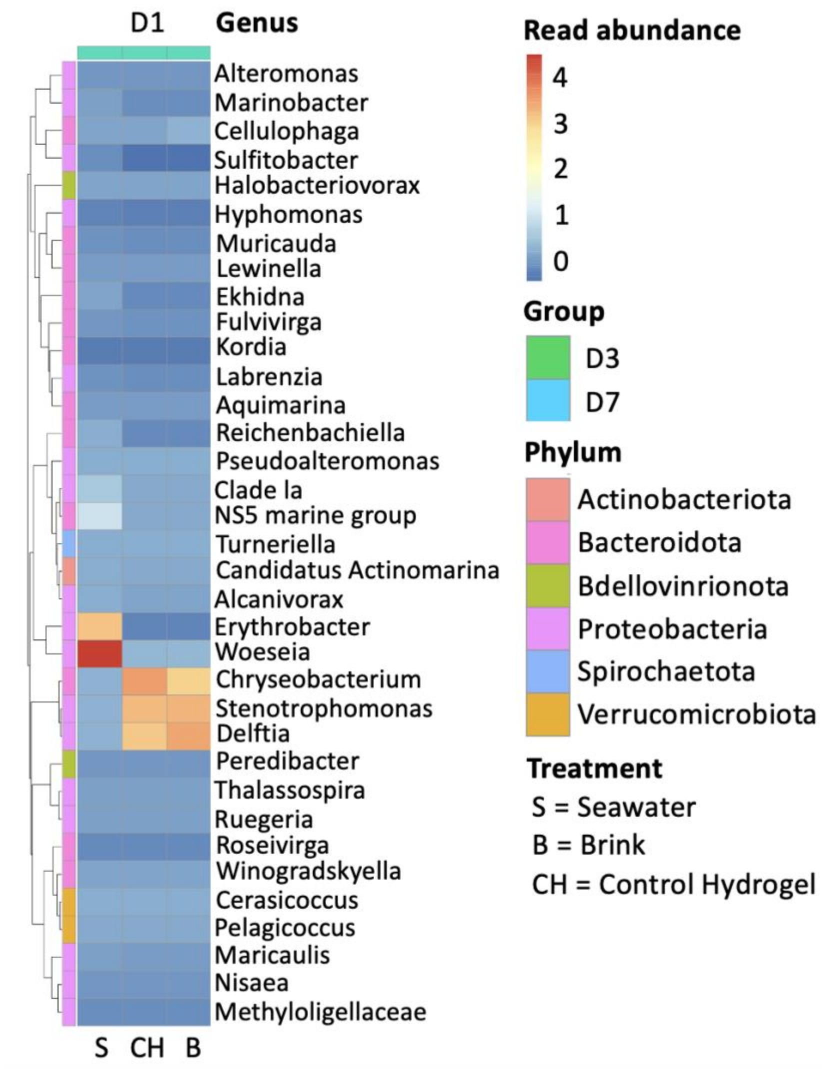


**Figure 5. Normalized read abundances of top 35 bacteria genera from 6 phyla associated with all samples: Seawater (S), Brink-coated plugs (B), and Control hydrogel plugs (CH).** Genera-level community analysis from the initial baseline sampling from day 1 (D1) of all treatments (*n*= 1) before the experiment started. Sample names on the x-axis, and y-axis represent the bacterial genera. Scale bars and cell colors represent the value of the distance between the raw score and the mean of the standard deviation, and the taxa list of genera is clustered by similarity.

**Coral larvae maintenance**

Fifteen colonies of *Pocillopora acuta* were retrieved from the fringing reef around Moku o Loʻe (Coconut Island), surrounding the Hawaiʻi Institute of Marine Biology (HIMB) on June 10th, 2023, immediately before their expected peak in planulation following the lunar 3rd quarter. After retrieval, they were transferred to a clean mesocosm tank (0.55 × 1.1 × 0.2 m, ~120 L), with constant flow-through of unfiltered seawater (~2 L $min^{-1}$), drawn from the adjacent reef, and adult corals were kept under a 30% shade with a neutral density cloth (max. irradiance of 1730 µmol photons $m^{-2}s^{-1}$) at HIMB. Planulae were released from the adult corals overnight and overflowed through the drain in the mesocosm tank. The overflowing seawater, containing the planulae, was directed to a larval collection bin (0.35 × 0.3 × 0.15 m, ~16 L). The seawater outflow for the larval collection bin was a 5 × 6 cm strainer covered by 100 µm mesh, to retain the larvae within the collection container. Each morning larvae were collected by pipette and transferred to 1 L conical larval rearing containers where they were kept in suspension under gentle aeration in unfiltered seawater. Larvae collected from the 11-13th of June were maintained in a single larval rearing container, whereas larvae collected from the 14-15th of June were set up in a second container, given the large number of larvae collected. The larval-rearing containers were maintained in a climate-controlled room (~23-25°C), and they received incident irradiance from a nearby window (max. irradiance ~200 µmol photons $m^{-2}s^{-1}$), where they were maintained before being added to settlement experiments.

Wild *Montipora capitata* gametes were retrieved from a local spawning slick in Kāneʻohe Bay near reef 11 (21.449248, - 157.796328) around the new moon in July 2023. The collection of gametes occurred from a patch reef, roughly several hundred square meters, which predominantly consists of *M. capitata* colonies comprising hundreds of parental genotypes. *M. capitata* is a hermaphroditic broadcast spawner that releases gametes at specific times. After collection, gamete bundles were brought back to HIMB, where they gradually separated and were fertilized in 50 mL conical tubes, rinsed, and added to filtered seawater conical rearing tanks, according to established best practices for *M. capitata* husbandry (Hancock et al., 2021). At 4 days post-fertilization, corals were reared to competency following established best practices (Rahnke et al., 2022) before being introduced to settlement experiments.
